# Remodeled Connexin 43 hemichannels alter cardiac excitability and promote arrhythmias

**DOI:** 10.1101/2022.03.08.483464

**Authors:** Mauricio A. Lillo, Manuel F. Muñoz, Kelli Gaul-Muller, Natalia Shirokova, Lai-Hua Xie, Diego Fraidenraich, Jorge E. Contreras

## Abstract

Connexin-43 (Cx43) is the most abundant protein forming gap junction channels (GJCs) in cardiac ventricles. In multiple cardiac pathologies, including hypertrophy and heart failure, Cx43 is found remodeled at the lateral side of the intercalated discs of ventricular cardiomyocytes. Remodeling of Cx43 has been long linked to spontaneous ventricular arrhythmia, yet the mechanisms by which arrhythmias develop are still debated. Using a model of a dystrophic cardiomyopathy, we previously showed that remodeled Cx43 function as aberrant hemichannels (non-forming GJCs) that alter cardiomyocyte excitability and, consequently, promote arrhythmias. Here, we aim to evaluate if opening of remodeled Cx43 can serve as a general mechanism to alter cardiac excitability independent of cellular dysfunction associated with a particular cardiomyopathy. To address this issue, we used a genetically modified Cx43 knock-in mouse (S3A) that promotes cardiac remodeling of Cx43 protein without apparent cardiac dysfunction. Importantly, when S3A mice were subjected to cardiac stress using the β-adrenergic agonist isoproterenol (Iso), they displayed acute and severe arrhythmias, which were not observed in WT mice. Pre-treatment of S3A mice with the Cx43 hemichannel blocker, Gap19, prevented Iso-induced abnormal electrocardiographic behavior. At the cellular level, when compared with WT, Iso-treated S3A cardiomyocytes showed increased membrane permeability and greater plasma membrane depolarization, which subsequently leads to triggered activity. These cellular dysfunctions were also prevented by Cx43 hemichannel blockers. Our results support the notion that opening of remodeled Cx43 hemichannels, regardless of the type of cardiomyopathy, is sufficient to mediate cardiac stress-induced arrhythmogenicity.

## INTRODUCTION

The intercalated discs of healthy cardiomyocytes contain gap junction channels, which act as low resistance channels to apposing cardiomyocytes(Kleber and Saffitz, 2014) and are essential for normal cardiac conduction. The connexin 43 (Cx43) gap junction forming protein is the most abundant connexin in the heart and is found in the working myocardium of the atrium and ventricle as well as the more distal regions of the Purkinje network (Gutstein et al., 2001; Oxford et al., 2007; Wang et al., 2013; Kleber and Saffitz, 2014; Gonzalez et al., 2018; Kim et al., 2019; Himelman et al., 2020). Multiple inherited or acquired cardiomyopathies, including dystrophic muscle dysfunction, arrhythmogenic right ventricular cardiomyopathy (ARVC), ischemia/reperfusion and hypertension, show an abnormal expression and remodeling of Cx43 (Severs et al., 2004a; Severs et al., 2004b; Severs et al., 2006; Oxford et al., 2007; Gonzalez et al., 2015; Kim et al., 2019). This dysregulation is thought to play a meaningful mechanistic role in the evolution of lethal cardiac arrhythmias (Severs et al., 2004a; Severs et al., 2004b; Severs et al., 2006; Gonzalez et al., 2015; Kim et al., 2019; Lillo et al., 2019; Himelman et al., 2020), likely by affecting the appropriate generation and spread of cardiac action potentials(Severs et al., 2006; Remo et al., 2011; Remo et al., 2012)

A common pathological feature of Cx43 cardiac remodeling is the dephosphorylation of a triplet of serine residues S325/S328/S330 at the carboxyl terminal domain of Cx43. These residues are usually phosphorylated by Ca^2+^/calmodulin protein kinase II (CaMKII) and/or casein kinase 1δ (CK1δ), which is a critical step for proper Cx43 localization at the intercalated discs (Qu et al., 2009; Huang et al., 2011; Remo et al., 2011). This mechanism of remodeling is strongly supported by data from Cx43 knock-in mouse lines where the three serine residues (S325/S328/S330) were replaced by phosphomimetic glutamic acid (S3E) or by non-phosphorylatable alanines (S3A). The S3E mice subjected to chronic pressure-overload hypertrophy were resistant to pathological Cx43 remodeling and to the induction of ventricular arrhythmias(Remo et al., 2011). Similarly, we showed that dystrophic mice containing S3E residues (mdxS3E) displayed absence of Cx43 remodeling with improved intracellular Ca^2+^ signaling and ROS production (Himelman et al., 2020). In addition, mdxS3E mice were less susceptible to arrhythmias and to the development of cardiomyopathy (Himelman et al., 2020). On the other hand, the S3A mice showed reduced gap junction formation and increased Cx43 remodeling in hearts subjected to chronic pressure-overload hypertrophy, consequently enhancing cardiac pathology (Remo et al., 2012; Himelman et al., 2020). Similar results were obtained for dystrophic mdxS3A mice. It was also shown that isolated hearts of S3A mice were more susceptible to inducible ventricular arrhythmias (Remo et al., 2011; Remo et al., 2012). Yet, the mechanisms by which they developed arrhythmias remain unclear (Remo et al., 2011; Remo et al., 2012).

Our group and others have recently proposed that remodeled Cx43 forms unpaired hemichannels (non-junctional channels) in ischemia and dystrophic mouse models (Wang et al., 2013; Gonzalez et al., 2018; Lillo et al., 2019; Himelman et al., 2020). Normally, unpaired Cx43 hemichannels are highly regulated to remain closed at the plasma membrane. However, we showed that mislocalized (lateralized) hemichannels in dystrophic mice are aberrantly activated upon cardiac stress leading to arrhythmias and sudden death (Gonzalez et al., 2015; Gonzalez et al., 2018). We found that β-adrenergic stimulation induced ventricular arrhythmias in dystrophic mouse models (mdx mice) but not in WT mice. This effect is prevented by injecting connexin mimetic peptides that selectively block Cx43 hemichannels but not GJ channels (Wang et al., 2013; Abudara et al., 2014; Gonzalez et al., 2015) or by genetically lowering levels of Cx43 (mdx:Cx43^+/−^)(Gonzalez et al., 2018). Similarly, in an ARVC mouse model where Cx43 is remodeled (Oxford et al., 2007; Kim et al., 2019), it has been shown that Cx43 hemichannels disrupt calcium homeostasis, promoting letal arrhythmogenesis (Kim et al., 2019). We discovered that β-adrenergic stimulation increases membrane permeability, depolarizes the plasma membrane and promotes triggered activity (TA) in dystrophic isolated cardiomyocytes via the opening of S-nitrosylated lateralized Cx43 hemichannels (Lillo et al., 2019). While the evidence for remodeled Cx43 hemichannels in cardiac dysfunction is emerging, it is unknown whether hemichannel opening is associated with unique cellular dysfunctions characteristic of a specific cardiomyopathy. For example, arrhythmias and sudden death are seen in mouse models of Duchenne muscular dystrophy, however, dystrophin loss and mechanical stress are present in addition to Cx43 remodeling.

Since Cx43 remodeling is observed in multiple cardiomyopathies, it is critical to explore whether opening of Cx43 hemichannels and consequent alterations in membrane excitability serve as a general mechanism of cardiac dysfunction independently of the etiology of the cardiomyopathy. To address this issue, we took advantage of the S3A mouse line, in which Cx43 remodeling is a result of direct genetic modification of the Cx43 protein rather than a consequence of cardiomyocyte pathological dysfunction. Upon cardiac stress, S3A mice developed acute ventricular arrhythmias consistent with previous observations in isolated hearts(Remo et al., 2011). In isolated S3A cardiomyocytes, β-adrenergic stimulation promoted membrane plasma depolarization and, subsequently, triggered activity (TA). These effects were prevented by blocking Cx43 hemichannels. Our data showed that opening of remodeled Cx43 hemichannels is sufficient to promote cardiac stress-induced arrhythmias.

## RESULTS

### S3A knock-in mice displayed remodeling of Cx43 hemichannels

We reassessed the distribution of Cx43 in S3A knock-in mouse hearts. Previously, it was shown that there is a reduction of Cx43 in the intercalated discs of S3A mice(Remo et al., 2011; Remo et al., 2012), however, Cx43 hemichannel lateralization in the absence of a cardiac pathological stimuli was not previously evaluated. To determine the distribution of Cx43 in wild-type and S3A hearts, we performed Cx43 immunohistochemistry in ventricular cryosections. Sections from 4- to 6-month-old mice were stained against Cx43 (green) in conjunction with the intercalated disc marker, N-cadherin (red). Confocal immunofluorescence imaging revealed that Cx43 signal in wild-type cardiomyocytes is prominently confined at the intercalated discs, overlapping with N-cadherin (Figure 1A). Conversely, in S3A hearts a substantial amount of Cx43 is found at lateral side of cardiomyocytes, where it does not overlap with N-cadherin. Quantification of relative Cx43 signal at the ID regions revealed that wild-type mice display significantly higher levels of Cx43 compared to S3A hearts (p <0.05, Figure 1A).

**Figure 1.**
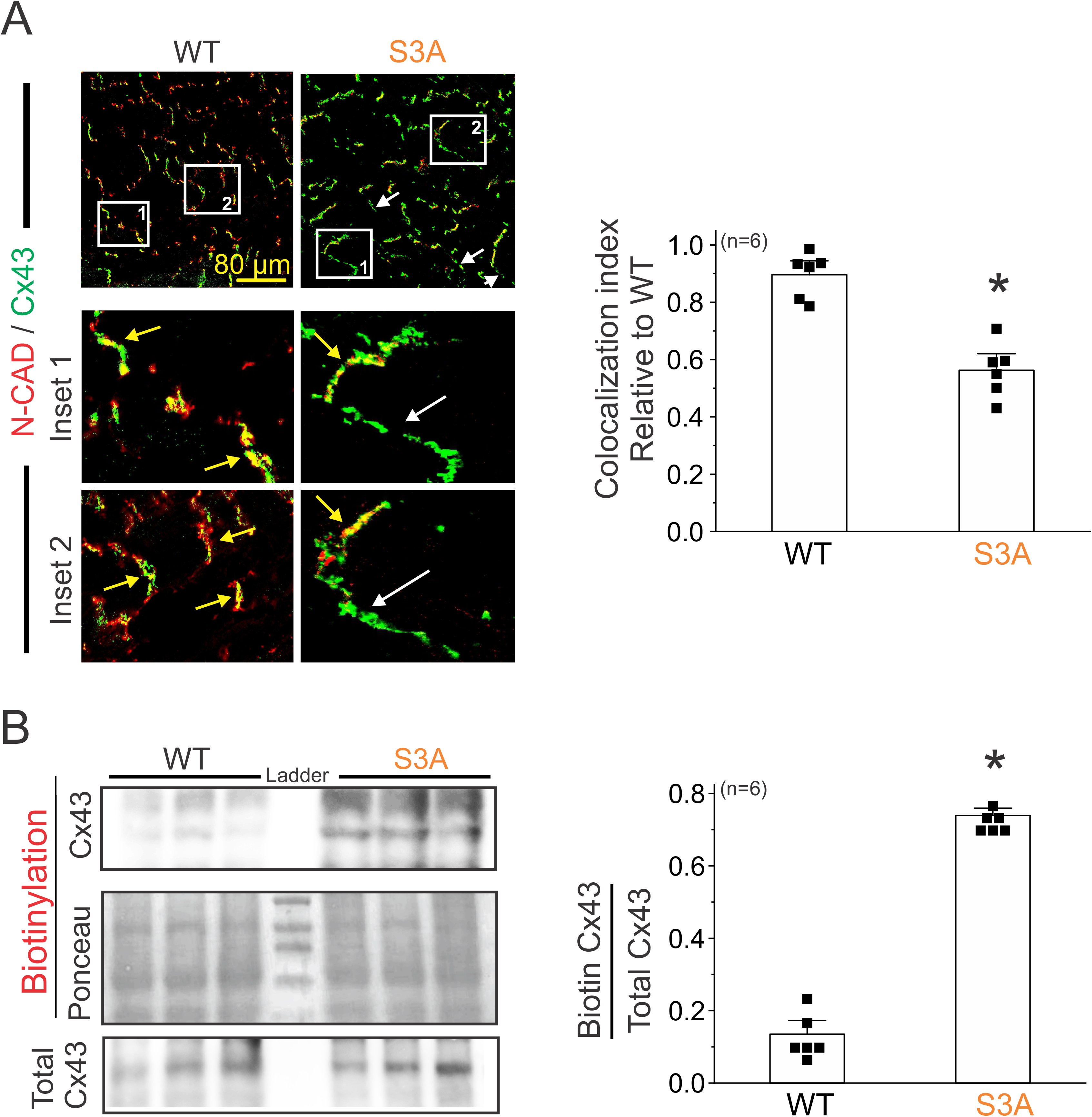
Cx43 hemichannels are lateralized in cardiomyocytes of S3A mice. **A)** Representative confocal immunofluorescence images of cardiac intercalated discs and lateral regions of cardiomyocytes stained with Cx43 (green) and N-Cadherin (red). Yellow and white arrows indicate intercalated disc and lateral side, respectively. Right: Quantification of Cx43/N-Cadherin co-localization in confocal immunofluorescence images. All data points were normalized to the WT group mean. Between Three and five images containing 15-20 IDs were analyzed per heart. Each dot represents the mean value for each mouse. Comparisons between groups were made using Student’s t test, *P<0.05 vs WT. **B**) Western blot analysis (left) and quantification (right) of Cx43 from biotin perfused hearts (biotinylation). Bottom row represents Cx43-immunoblotted samples from heart lysates prior to pulldown (total Cx43). Biotinylated Cx43 levels were expressed as fold change relative to total Cx43 protein levels per sample. The number in parentheses indicates the *n* value. Comparisons between groups were made using Student’s t test, *P<0.05 vs WT.

Next, we attempted to compare levels of lateralized Cx43 in cardiomyocytes from S3A versus wild-type hearts. To achieve this, we perfused isolated hearts from wild-type and S3A mice with cell-impermeable biotin (MW:244.31), which strongly binds to non-junctional Cx43 hemichannels as we have previously demonstrated (Lillo et al., 2019; Himelman et al., 2020). After aorta cannulation and biotin perfusion, hearts were homogenized, biotinylated proteins were pulled down with streptavidin beads, run on SDS PAGE and probed for Cx43. Western blot analysis of the biotinylated fraction showed that S3A hearts have nearly 8-fold higher Cx43 protein levels at the lateral region compared to wild-type hearts. (Figure 1B). Both immunofluorescence and biochemical studies confirm that S3A cardiomyocytes display Cx43 remodeling as reported for various cardiac pathologies.

### Iso-induced cardiac arrhythmias in S3A mice were prevented by blocking Cx43 hemichannels

We have previously shown that remodeled Cx43 protein mediates cardiac stress-induced arrhythmias in dystrophic mice via opening of Cx43 hemichannels(Gonzalez et al., 2018; Lillo et al., 2019; Himelman et al., 2020). Thus, we tested whether Cx43 remodeling in S3A mice leads to susceptibility to arrhythmias via a similar mechanism. We performed whole animal electrocardiograms (ECGs) using a telemetry system before and after isoproterenol (Iso) challenge in 4- to 6-month-old wild-type and S3A mice under control condition or after pre-treatment with the Cx43 hemichannel inhibitor, Gap19 (Wang et al., 2013; Abudara et al., 2014) (10 μg/kg via retro-orbital injection). Representative ECGs recorded from wild-type and S3A mice treated with Gap19 are shown in Figure 2A. None of the mice in each group presented arrhythmias under baseline conditions. Following Iso administration, S3A mice developed severe arrhythmias, including premature ventricular contractions (PVCs), ventricular tachycardia (VT), and atrioventricular (AV) block (Figure 2A and 2B). In contrast, when pre-treated with the Cx43 hemichannel blocker, Gap19, S3A mice displayed no abnormalities or very infrequent PVCs within 1 hour post-Iso injection (Figure 2A and 2B). We further monitored arrhythmogenic behavior during the 24 hours following Iso stimulation in both wild-type and S3A mice. The S3A mice displayed increased arrhythmogenic events compared to wild-type mice within 24 hours post-Iso injection (130.4 ± 4.5 and 10.2± 1.5, respectively) (Figure 2C). The abnormal arrhythmogenic behavior detected in S3A mice was reduced following pretreatment with Gap19 (26.3 ± 6.2) (Figure 2C).

**Figure 2.**
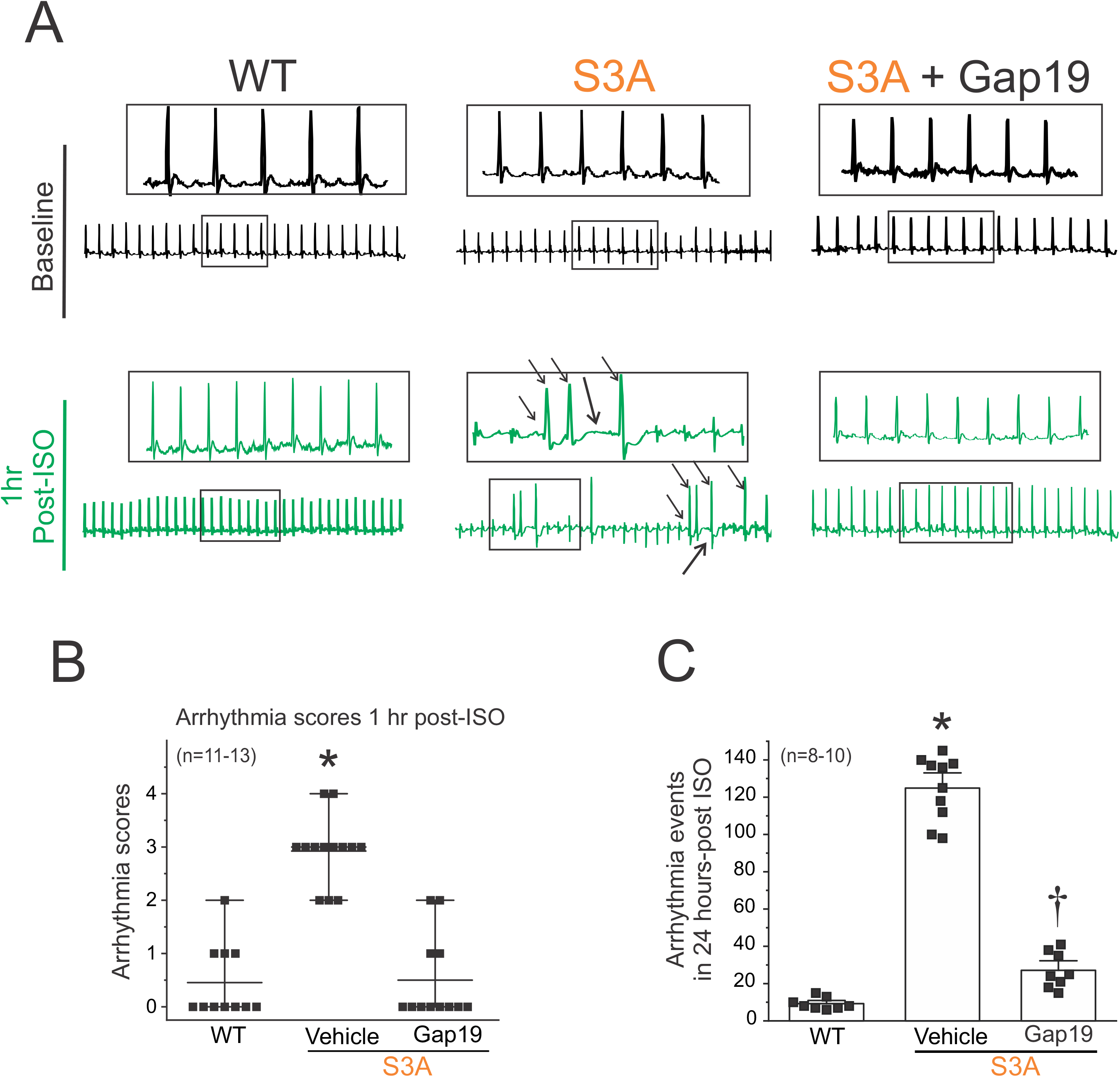
β-adrenergic cardiac stress promotes arrhythmogenic phenotypes in S3A mice, which are prevented by blocking Cx43 hemichannels. **A**) Representative ECG traces obtained from 4–6 month-old mice using an *in vivo* telemetry system in WT, S3A and S3A mice treated with Gap19 via retroorbital injection (10 μg/kg). Scale bar is 100ms for all traces. Black arrows indicate arrhythmia events. **B)** Arrhythmia scores 1 hour post-Isoproterenol (Iso) challenge based on pre-determined scale where 0 = no arrhythmias, 1 = single PVCs, 2 = double PVCs, 3 = triple PVCs or non-sustained VT, 4 = sustained VT or AV block, 5 = death. Comparisons between groups were made using two-way ANOVA plus Tukey post-hoc test, * p<0.0001 versus WT; † p<0.0001 versus S3A + Gap19. **C)** Quantification of arrhythmogenic events (including PVC, double PVC, VT, or AV block) during the 24 hours post-Iso challenge in WT, S3A and S3A mice treated with Gap19 via retroorbital injection (10 μg/kg). The number in parentheses indicates the *n* value. Comparisons between groups were made using two-way ANOVA plus Tukey post-hoc test. *<0.05 vs WT, †P<0.05 vs S3A.

### Iso-induced opening of Cx43 hemichannels led to increased membrane permeability, triggered activity and prolonged action potentials in S3A cardiomyocytes

We previously showed that opening of remodeled Cx43 hemichannels increased membrane excitability and evoked triggered activity in Duchenne dystrophic isolated cardiomyocytes upon treatment with isoproterenol (Lillo et al., 2019). Thus, we examined whether these pathological features are replicated in S3A cardiomyocytes. Figure 3A displays representative traces of cardiac action potentials (APs) from wild-type and S3A isolated cardiomyocytes induced by an injection of 2 nA current under current-clamp conditions. Electrical stimulation of S3A isolated cardiomyocytes showed a small increase in triggered activity (TA) compared to wild-type cardiac cells, 8.4 ± 0.4 and 1.8 ± 0.05 per minute, respectively (Figure 3A and 3B). Treatment of cells with 1μM Iso induced TA in S3A but not wild-type cardiomyocytes, 56.8 ± 3.8 and 2.4 ± 0.09 per minute, respectively (Figure 3A and 3B). To determine the role of Cx43 hemichannels in Iso-induced TA, we tested whether the addition of two different Cx43 hemichannel-specific blockers, Gap19 and Cx43 antibody (Gonzalez et al., 2015; Lillo et al., 2019), into the patch pipette reduced TA. Iso-induced TA was significantly reduced (by > 70 %) in S3A cardiomyocytes treated with both Gap19 peptide (10.1 ± 2.1 per minute) or AbCx43 (6.4 ± 0.81 per minute) (Figure 3A and 3B).

**Figure 3.**
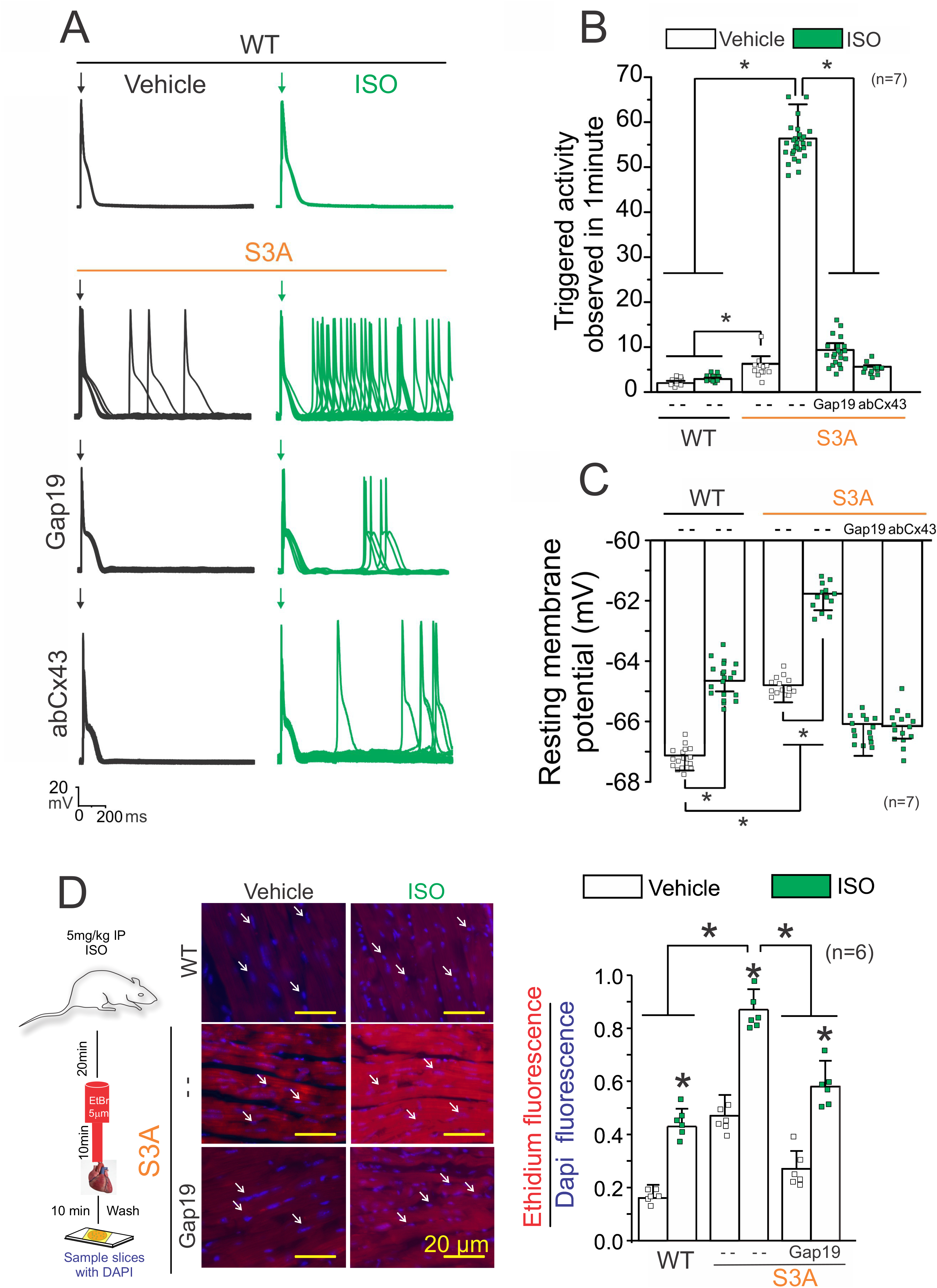
Isoproterenol induces TA in S3A cardiomyocytes via opening of Cx43 hemichannels. **A)** Representative action potential traces of WT and S3A isolated cardiomyocytes. Cells were stimulated with 1μM isoproterenol (Iso, green) in the absence or presence of Cx43 blockers contained inside the pipette: Gap19 (232ng/μL) and Cx43 antibody (abCx43; 2.5 ng/μL). Arrow indicates electrical stimulation pulse. **B)** Quantification of TA observed in **(A)**. Comparisons between groups were made using two-way ANOVA plus Tukey post-hoc test, *P<0.05. **C)** Resting membrane potential of WT and S3A cardiomyocytes. The number in parentheses indicates the *n* value. Comparisons between groups were made using two-way ANOVA plus Tukey post-hoc test, *P<0.05. **D**) Evaluation of Cx43 hemichannel activity in the whole heart via ethidium uptake. Isolated hearts were perfused with buffer containing 5μM ethidium after vehicle or Iso (5mg/kg, IP). The number in parentheses indicates the *n* value. Comparisons between groups were made using two-way ANOVA plus Tukey post-hoc test. *P<0.05.

We next tested whether Iso-induced TA in S3A cardiomyocytes is correlated with alterations in the resting membrane potential (*Vm*), which in turn is produced by the distorted lateralized Cx43 hemichannels. Figure 3C shows that S3A cardiomyocytes are more depolarized with respect to wild-type cardiomyocytes under resting conditions, with *Vm* values of −64.1 ± 1.8 mV and −67.2 ± 2.1 mV, respectively. Furthermore, Iso stimulation depolarized both S3A and wild-type cardiomyocytes to *Vm* values of −61.7 ± 2.1 mV and −64.3 ± 1.2 mV, respectively (Figure 3C).

Remarkably, when Cx43 hemichannels blockers (Gap19 or AbCx43) were included in the pipette solution, the *Vm* in S3A cardiomyocytes was restored to values similar to those observed in wild-type cardiac cells (Figure 3C). These results support that plasma membrane depolarization in S3A cardiomyocytes is caused by the disrupted activity of Cx43 hemichannels.

Uptake from the extracellular space of hemichannel-permeable fluorescent molecules, such as ethidium bromide (EtBr), is generally used to identify opening of Cx43 hemichannels (Contreras et al., 2003; Figueroa et al., 2013; Johnson et al., 2016). To confirm that Cx43 lateralized hemichannels are functional in S3A cardiomyocytes, we applied a semi-quantitative *in-situ* assay utilizing perfused isolated hearts as previously reported (Lillo et al., 2019; Himelman et al., 2020). Under normal conditions, S3A hearts showed about five-fold greater EtBr uptake than wild-type hearts (Figure 3D). Iso stimulation enhanced EtBr uptake in both genotypes, but a significantly larger uptake was detected in S3A hearts (Figure 3D). *In vivo* treatment with Gap19 via retro-orbital injection prior to Iso administration (5mg/Kg, IP) significantly reduced dye uptake in S3A hearts under both normal and Iso stimulated conditions (Figure 3D).

Previous work has showed a significant increase in action potential duration (APD) dispersion in S3A mice compared to wild-type (Remo et al., 2011). Thus, we analyzed whether specifically Cx43 hemichannels could alter APD interval in isolated cardiomyocytes from wild-type and S3A mice, before and after treatment with 1μM Iso. Consistent with previous findings (Lillo et al., 2019), S3A cardiomyocytes displayed longer APD with respect to wild-type cardiomyocytes under vehicle conditions, with APD values of 133.28 ± 3.3 milliseconds (ms) and 104.8 ± 1.4 ms, respectively (Figure 4A and 4B). Furthermore, Iso stimulation increased both S3A and WT cardiomyocytes to APD values of 180.6 ± 3.3 ms and 134.83 ± 1.86 ms, respectively. When Cx43 hemichannels blockers (Gap19 or AbCx43) were contained in the pipette solution, APD in S3A cardiomyocytes was restored to values similar to those observed in wild-type cardiac cells (Figure 4B). These results indicate that activity of Cx43 hemichannels influences action potential duration in S3A cardiomyocytes. The fact that remodeled Cx43 hemichannels significantly alter excitatory properties of cardiomyocytes is consistent with a role in conferring the pro-arrhythmic phenotype observed in these mice.

**Figure 4.**
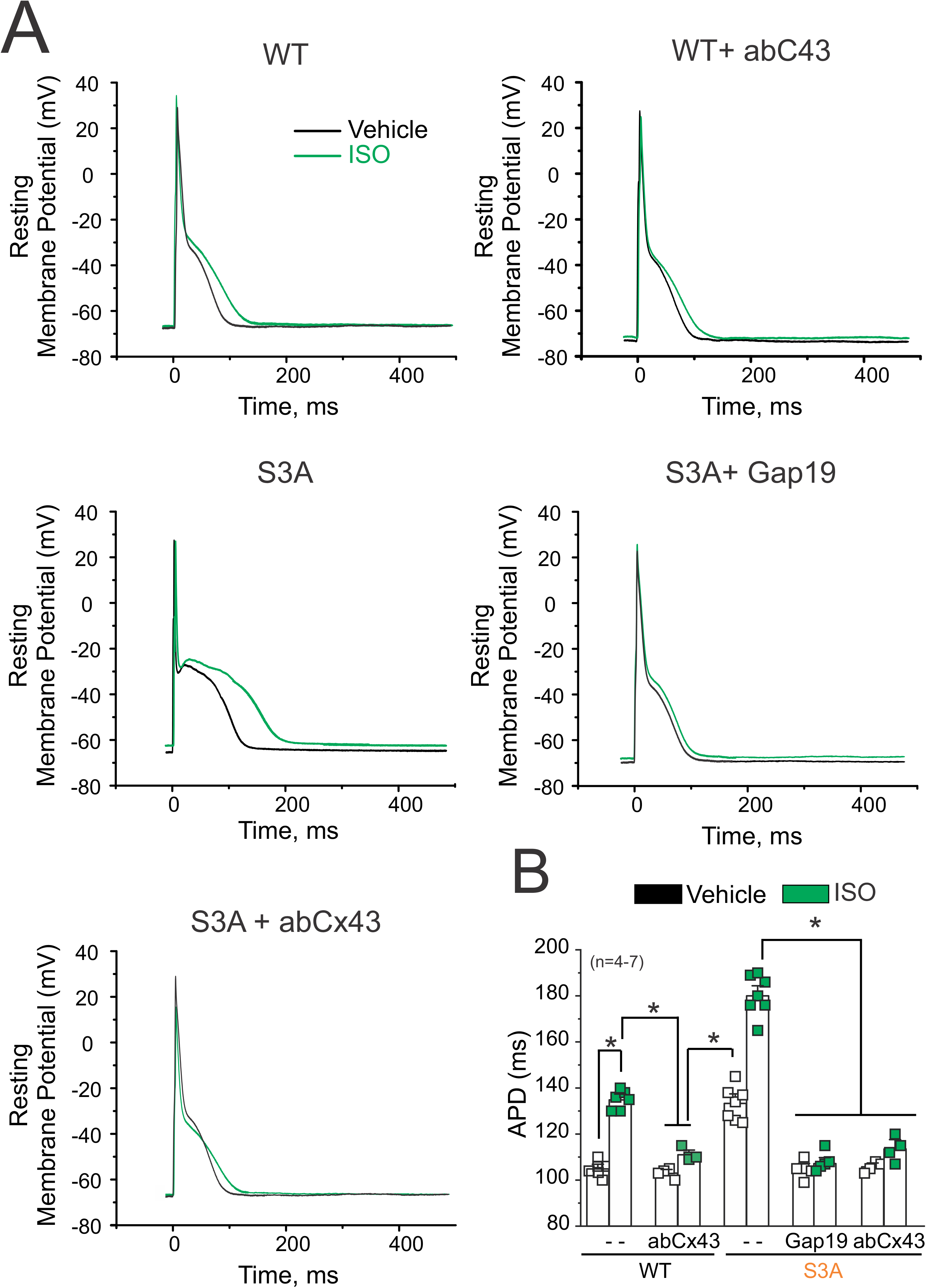
Cx43 hemichannels are involved in action potential duration (APD) prolongation in WT and S3A isolated cardiomyocytes. **A)** Representative action potential traces of WT and S3A isolated cardiomyocytes. Cells were stimulated with 1μM isoproterenol (Iso, green) in the absence or presence of Cx43 blockers contained inside the pipette: Gap19 (232ng/μL) and Cx43 CT antibody (abCx43; 2.5 ng/μL). **B)** Quantification of APD observed in **(A)**. The number in parentheses indicates the *n* value. Comparisons between groups were made using two-way ANOVA plus Tukey post-hoc test, *P<0.05.

### β-adrenergic cardiac stress evokes triggered activity in S3A cardiac cells via S-nitrosylation of lateralized Cx43 hemichannels

We recently reported that S-nitrosylation of Cx43 hemichannels in the dystrophic cardiomyocytes is necessary to promote Iso-induced disruption of membrane excitability (Lillo et al., 2019). Thus, we evaluated whether Iso-induced TA in S3A cardiomyocytes is mediated by NO production and S-nitrosylation of remodeled Cx43. Figure 5A shows representative action potentials in S3A isolated cardiomyocytes that were evoked by electrical stimulation in the presence of 100 μM L-NAME, a non-selective NOS inhibitor (Pfeiffer et al., 1996). L-NAME treatment greatly reduced the incidence of Iso-evoked TAs in S3A cardiomyocytes (6.7 ± 2.1 per minute, Figure 5B). Consistently, L-NAME treatment significantly reduced the Iso-induced decrease in resting membrane potential of S3A cardiomyocytes (Figure 5C) and restored it to similar values observed in wild-type cardiomyocytes (−67.2 ± 2.1 Figure 4C). To validate a direct role of NO on the generation of TA in S3A cardiomyocytes, we directly stimulated S3A cardiomyocytes with a NO donor, sodium 2-(N, N-diethylamino)-diazenolate-2-oxide (DEENO, 1μM). While some TA was observed in S3A isolated cardiomyocytes treated with vehicle (8.5 ± 2.4 per min), there were significantly more TA events in S3A cardiomyocytes when treated with DEENO (61.3 ± 5.1 per minute; Figure 5A and 5B). In addition, exogenous NO application depolarized the membrane S3A cardiomyocytes to *Vm* of −61.5 ± 2.6 (Figure 5C). These results support the notion that Cx43 hemichannels mediate aberrant electrical activity in S3A cardiomyocytes via S-nitrosylation of Cx43 proteins.

**Figure 5.**
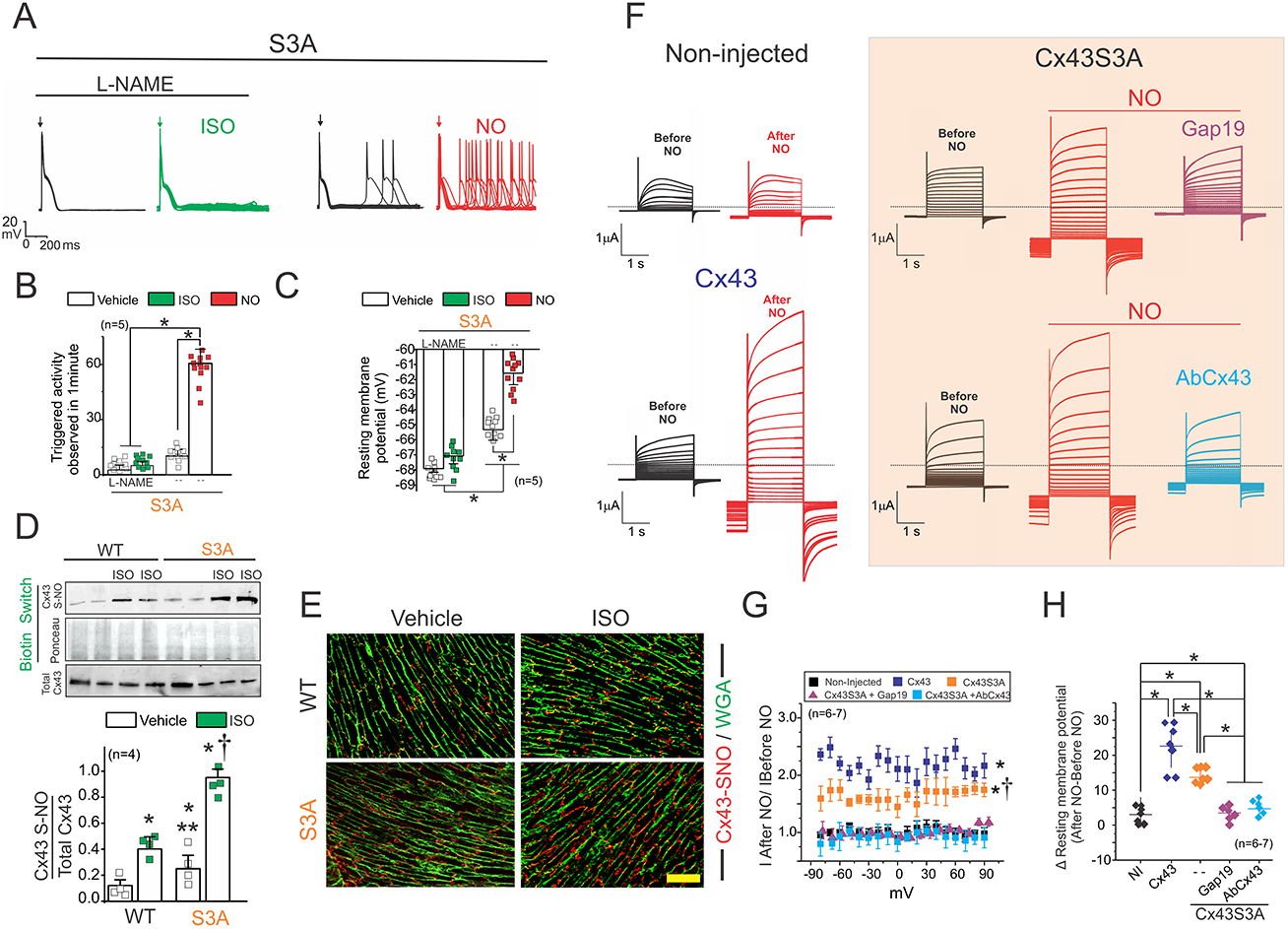
β-adrenergic cardiac stress evokes triggered activity in S3A cardiac cells via S-nitrosylation of lateralized Cx43 hemichannels. **A)** Representative action potential traces of S3A isolated cardiomyocytes. Cells were stimulated with 1μM isoproterenol (Iso) in the presence of 100μM L-NAME (NOS blocker) or 1 μM DEENO (NO donor). Arrow indicates electrical stimulation. **B)** Quantification of TA induced by Iso observed in **(A)**. The number in parentheses indicates the *n* value. Comparisons between groups were made using two-way ANOVA plus Tukey post-hoc test *P<0.05. **C)** Resting membrane potential of S3A cardiomyocytes. Comparisons between groups were made using two-way ANOVA plus Tukey post-hoc test. **D**) Top and middle gels were loaded with S-nitrosylated proteins pulled down from heart samples using the biotin switch assay. Top gel was then blotted against Cx43. The middle gel is the corresponding ponceau staining. Lower blot was loaded using total cardiac proteins and blotted against Cx43. The bottom graph is the quantification for 2 independent blots using the ratio for SNO-Cx43/Total Cx43. The number in parentheses indicates the *n* value. Comparisons between groups were made using two-way ANOVA plus Tukey post-hoc test. *P<0.05 vs WT control, **P<0.05 vs WT ISO, †<0.05 vs WT Iso. **E)** Analysis performed by Proximity Ligation assay (PLA) of the interaction between Cx43 and S-nitrosylation. Plasma membrane stained with wheat germ agglutinin (WGA) and S-nitrosylated Cx43 (Cx43-SNO) are shown in green and red, respectively. Representative images of n = 3 per group. **F)** Representative current traces before and after application of 10 μM DEENO in a non-injected oocyte or an oocyte expressing Cx43 and Cx43S3A. Oocytes were clamped to −80 mV, and square pulses from −80 mV to +90 mV (in 10 mV steps) were then applied for 2s. At the end of each pulse, the membrane potential was returned to −80 mV. Intracellular injection of Gap19 (232 ng/μL) or a Cx43 CT antibody (2.5 ng/μL) reduces NO-induced Cx43 hemichannel currents. **G)** Normalized currents were obtained from the ratio between recorded current after and before DEENO treatment. The number in parentheses indicates the *n* value. Comparisons between groups were made using two-way ANOVA plus Tukey post-hoc test, *P<0.05 vs Non-Injected, †<0.05 vs Cx43. **H**) Changes in resting membrane potential in the presence or absence 10 μM DEENO. Cx43 hemichannel blockers restore normal resting membrane potential. Comparisons between groups were made using one-way ANOVA, *P<0.05.

Next, we used the biotin switch assay to compare Cx43 S-nitrosylation levels in wild-type versus S3A hearts. We found that levels of S-nitrosylated Cx43 in S3A cardiac tissue is near 2-fold greater than in the wild-type heart under control conditions (Figure 5D). *In vivo* Iso treatment resulted in an increase in levels of S-nitrosylated Cx43 in both wild-type and S3A hearts by about to 2- and 4-fold when compared to their respective controls (Figure 5D). To analyze the subcellular localization of Cx43 S-nitrosylated proteins, we used the Proximity Ligation Assay (PLA) using antibodies that recognize S-nitrosylated proteins (S-NO) and the C-terminus of Cx43, as we have previously described (Lillo et al., 2019). We detected Cx43-NO signals (red stain) in the intercalated disc in both WT and S3A cardiac tissue under vehicle conditions (Figure 5E). However, after treatment with Iso, S-nitrosylated Cx43 was only found at the lateral side of cardiomyocytes in S3A hearts (Figure 5E).

To confirm that S3A mutations do not alter NO response and biophysical properties of Cx43, we assessed Cx43S3A hemichannel currents in *Xenopus* oocytes using the two-electrode voltage clamp technique. As we previously reported (Lillo et al., 2019), in the absence of NO donors the voltage-activated leak currents in Cx43 expressing oocytes are similar to those observed in non-injected oocytes. However, application of 10μM DEENO evoked an increase in plasma membrane conductance at positive and negative voltages in Cx43 expressing oocytes, but not in non-injected oocytes (Figure 5F). Oocytes expressing Cx43S3A mutations also displayed an increase in leak currents in response to DEENO (Figure 5F orange box). NO-induced leak currents in Cx43S3A expressing oocytes were reduced in oocytes that were intracellularly injected with Cx43 hemichannel blockers, Gap19 or AbCx43 (Figure 5F orange box). The increase in NO-induced leak currents was 2- and 1.6-fold in Cx43 and Cx43S3A expressing oocytes, respectively (Figure 5F and 5G). In addition, we found that NO evoked a more depolarized *Vm* in oocytes expressing Cx43 and Cx43S3A when compare with non-injected oocytes (Figure 5H). Inhibition of Cx43 hemichannels with Gap19 or AbCx43 prevented NO-induced *Vm* depolarization in Cx43S3A expressing oocytes (Figure 5H). Our results confirm that Cx43S3A hemichannels are activated by NO and result in resting membrane depolarization in *Xenopus* oocytes, which is similar to what was reported above for isolated S3A cardiomyocytes.

## DISCUSION

Cx43 remodeling is well accepted as a feature of cardiac pathologies involving spontaneous ventricular arrhythmia. In prior studies, we described that remodeled Cx43 forms undocked connexin hemichannels, whose activity leads to arrhythmias and cardiac dysfunction in a mouse model of Duchenne Muscular Dystrophy (Gonzalez et al., 2018; Lillo et al., 2019; Himelman et al., 2020). Here we examine whether opening of remodeled Cx43 hemichannels serves as a general mechanism underlying dysfunctional cardiomyocyte excitability or it is a pathological dependent phenomenon. Using the S3A knock-in mice with genetically engineered remodeled Cx43 proteins, we showed that Iso-induced cardiac stress promoted a substantial number of arrhythmogenic events that were prevented by blockade of Cx43 hemichannels. At the cellular level, we found that blockade of Cx43 hemichannels prevents abnormal changes in membrane permeability, plasma membrane depolarization and Iso-evoked TA. Our data are consistent with the idea that opening of Cx43 hemichannels is, at least, one of the mechanisms by which Cx43 protein remodeling is involved in the development of arrhythmias.

Cx43 remodeling is observed in pathologic studies of human hearts with a broad class of collected arrhythmic symptoms, including ischemic and hypertrophic cardiomyopathies, acquired diseases such as arrhythmogenic right ventricular cardiomyopathy (ARVC), genetic diseases associated with somatic or germline mutations in connexin genes, and muscular dystrophic cardiomyopathy (Gutstein et al., 2001; Gollob, 2006; Bruce et al., 2008; Saffitz, 2009; Thibodeau et al., 2010; Remo et al., 2012; Gonzalez et al., 2018; Kim et al., 2019; Lillo et al., 2019; Himelman et al., 2020). Mechanisms linking Cx43 remodeling to the enhanced propensity for arrhythmic activity have been associated with dysfunction of multiple cellular signaling pathways (Gollob, 2006; Saffitz et al., 2009; Thibodeau et al., 2010; Kim et al., 2019). Abnormal localization and/or gating of ion channels (including Nav1.5, NCX, K^+^ currents, among others) disturbs the highly organized temporal and spatial pattern of cardiac excitation (Gollob, 2006; Saffitz et al., 2009; Thibodeau et al., 2010; Kim et al., 2019). The resulting slow, dysfunctional and often heterogeneous conduction favors reentrant activity, thus promoting arrhythmias, fibrillation, heart failure or sudden death (Gollob, 2006; Saffitz et al., 2009; Thibodeau et al., 2010; Kim et al., 2019). However, among all these studies only few have considered the role of Cx43 hemichannels as a mediator of arrhythmogenic behavior (Kim et al., 2019; Lillo et al., 2019; Himelman et al., 2020; De Smet et al., 2021; Lillo and Contreras, 2021).

We have previously demonstrated that β-adrenergic stimulation in dystrophic cardiomyocytes evoked TA and calcium entry via opening S-nitrosylated Cx43 hemichannels (Gonzalez et al., 2018; Lillo et al., 2019; Himelman et al., 2020). TA and calcium entry were prevented by Cx43 hemichannel blockers (Lillo et al., 2019; Himelman et al., 2020) or by genetically reducing levels of Cx43 in a dystrophic mouse model(Lillo et al., 2019). They were also prevented in dystrophic cardiac cells where Cx43 was genetically modified to prevent remodeling (Himelman et al., 2020).

In line with these finding, a recent report showed that blockade of Cx43 hemichannels in ventricular cardiomyocytes from heart failure patients prevents cardiac stress induced spontaneous Ca^2+^ release, delayed after depolarizations (DADs) and TA (De Smet et al., 2021). In the present work, we showed that β-adrenergic stimulation results in depolarization of the cardiac plasma membrane in both WT and S3A cardiomyocytes. However, the depolarization of S3A cells was larger and evokes TA. Cx43 hemichannel blockers restored the resting membrane potential in Iso-treated S3A cardiomyocytes to levels observed in WT and significantly decreased TA. Similar to what was described by us in dystrophic cardiomyocytes(Lillo et al., 2019), Iso-treated S3A cardiomyocytes only produced a rightward shift of ~ 7 mV in the resting membrane potential compared with WT cells. We previously explained that severe changes in the resting membrane potential that would be caused by full opening of Cx43 hemichannels is prevented by physiological extracellular Ca^2+^ concentration, which significantly decreases Cx43 hemichannel open probability (Contreras et al., 2003; Lillo et al., 2019).

Arrhythmogenic events in S3A mice were also detected in the absence of β-adrenergic stimulation mainly during the dark phase of the light/dark cycle (Figure 4B and Figure S1), when mice were most active. Consistently, at the cellular level, we observed TA in isolated S3A cardiomyocytes without Iso-stimulation. Thus, both TA and arrhythmias are likely linked to sporadic increases of intracellular calcium in cardiomyocytes. It is well known that dysregulation of calcium handling promotes arrhythmogenic phenotypes in several animal models of heart failure (Gollob, 2006; Saffitz et al., 2009; Thibodeau et al., 2010; Kyrychenko et al., 2013; Gonzalez et al., 2018; Kim et al., 2019; Lillo et al., 2019; Himelman et al., 2020). Disruption of Ca^2+^ homeostasis leads to DADs, EADs, and TA. For instance, in plakophilin-2-2 (PKP2) deficient mice, Cx43 hemichannel activity is upregulated, resulting in increased intracellular Ca^2+^ levels and, consequently, arrhythmogenic behavior(Kim et al., 2019). In this work, the authors suggest that Cx43 hemichannels at the surface membrane of cardiomyocytes might offer a pathway for Ca^2+^ entry into cardiomyocytes from PKP2 deficient mice. Hence, blockade of Cx43 hemichannels with Gap19 or using PKP2 mice with lower levels of Cx43 (PKP2: Cx43^+/−^) leads to the rescue of Ca^2+^ homeostasis and reduction of membrane leakage to small molecules (evaluated by Lucifer yellow uptake) in isolated hearts (Kim et al., 2019). Our current data in hearts from S3A mice are in line with these previous findings. This idea is further supported by our ethidium uptake assays showing that Iso-treatment produced a substantial increase in membrane permeability of S3A hearts that is significantly reduced by the blockade of Cx43 hemichannels. We inferred that cardiac stress-induced TA results from enhanced activity of Cx43 hemichannels in S3A cardiomyocytes.

β-adrenergic or electrical cardiac stimulation activate cellular signaling pathways that include multiple enzymes such as Akt, PKA, Ca^2+^/calmodulin-dependent protein kinase II and nitric oxide synthases (Zhu et al., 2003; Grimm and Brown, 2010; Mani et al., 2010; Hegyi et al., 2019; Lillo et al., 2019). These pathways are responsible for the phosphorylation or S-nitrosylation of multiple cardiac ion channels (Vielma et al., 2016), including Cx43 proteins (Lillo et al., 2019). These posttranslational modifications modulate cardiac electrical activity and Ca^2+^ homeostasis (Zhu et al., 2003; Grimm and Brown, 2010; Thibodeau et al., 2010; Kyrychenko et al., 2013; Hegyi et al., 2019; Kim et al., 2019). Recent evidence indicates that cardiac Cx43 hemichannels under physiological conditions can open as a result of a moderate increase in intracellular Ca^2+^ directly following Iso-induced RyR2 activation (Lissoni et al., 2019). The data convincingly show a functional cross-talk between Cx43 and RyR2 receptors. In addition, Cx43 hemichannels were found interacting in microdomains with RyR2 at the ID, close to the perinexus. The authors reported that elevations of diastolic Ca^2+^ through caffeine or β-adrenergic-stimulation promoted activity of single Cx43 hemichannels in isolated and paired ventricular cardiomyocytes, in pig and mouse(De Smet et al., 2021). Interestingly, they also showed that further hemichannel opening evoked few DADs or TA in healthy cardiomyocytes.

Another feature that has been associated with arrhythmogenic behavior is the prolongation of action potential duration (APD) (Tse, 2016). Consistently with previous results (Remo et al., 2011; Lillo et al., 2019; De Smet et al., 2021; Lissoni et al., 2021), our data indicate that Cx43 hemichannels modulate the duration of the action potential in both WT cardiac cells and S3A. This observation was detected in other cardiac models, where Cx43 proteins are also remodeled. Ghazizadeh et al. reported, in an atrial fibrillation cardiac model, that inhibition of Cx43 hemichannels rescues the changes in APD and membrane leakage (Ghazizadeh et al., 2020). By blocking Cx43 hemichannel activity, we restored the APD in cells with Cx43 lateralized hemichannels, such as Dystrophic cardiac cells(Lillo et al., 2019) or S3A isolated cells, with and without isoproterenol stimulation (Figure 5). Altogether, these results strongly support the idea that Cx43 hemichannels modulate cardiac excitability in pathological cardiomyocytes and, perhaps, in healthy ones.

The RyR2 is one of the most studied Ca^2+^ channels in cardiac pathologies. Dysregulation in phosphorylation or nitrosylation patterns modifies RyR2 activity and affects Ca^2+^ handling (Gonzalez et al., 2007; Forrester et al., 2009; Fauconnier et al., 2010; Lima et al., 2010), changing electrical cardiac performance and inducing heart arrhythmias, heart failure, or even sudden death in several cardiac animal models (Marx et al., 2000; Oxford et al., 2007; Fauconnier et al., 2010; Grimm and Brown, 2010; Mani et al., 2010; Remo et al., 2012; Kim et al., 2019; Lillo et al., 2019; Himelman et al., 2020). In dystrophic mice, both RyR2 and Cx43 become hyper-nitrosylated upon β-adrenergic stimulation (Fauconnier et al., 2010; Lillo et al., 2019; Vielma et al., 2020). Inhibition of nitric oxide production by L-NAME reduced S-nitrosylation of Cx43 hemichannels in dystrophic mice as well as the related increase in membrane depolarization and the development of TA and arrhythmias (Lillo et al., 2019). Similarly, in the hearts of S3A mice Cx43 becomes highly S-nitrosylated in the lateral membrane of cardiomyocytes and L-NAME treatment of isolated S3A cardiomyocytes diminished Cx43 hemichannel-mediated membrane depolarization and TA. We also confirmed that Cx43 S3A hemichannels are activated by NO when expressed in heterologous *Xenopus* oocytes, although the activation was slightly less than Cx43 WT. Yet, activation of Cx43 S3A hemichannels was sufficient to produce large membrane depolarizations in *Xenopus* oocytes. As expected, the activation of Cx43 S3A hemichannels was significantly inhibited when Gap19 or AbCx43 was injected into the oocytes. These inhibitors also restore changes in oocyte membrane potentials mediated by Cx43 S3A hemichannels in a similar manner to that observed in S3A cardiomyocytes. These controls validate a role for the opening of Cx43 hemichannels mediating changes in membrane excitability.

Importantly, our findings in the S3A knock-in mice support a pathological role for remodeled Cx43 hemichannels, which alter membrane excitability and enhance cardiomyocyte dysfunction upon cardiac stress. These findings may help to develop therapeutic approaches for multiple cardiomyopathy phenotypes where Cx43 proteins are remodeled.

## MATERIALS AND METHODS

### Mouse Breeding and Genotyping

S3A founder mice colonies were a gift from Dr. Glenn Fishman (New York University). WT mice were purchased in Jackson Labs and examined at time points of 5-6 months. All animal investigations were supported by the IACUC of Rutgers New Jersey Medical School and UC Davis Medical School performed following the NIH guidelines.

### Cardiomyocytes

Ventricular myocytes were enzymatically isolated from WT or S3A mice. Mice were heparinized (5000 U/kg) and then anesthetized with overdosed isoflurane. The hearts were removed and perfused at 37 °C in Langendorff method with nominally Ca^2+^-free Tyrode’s solution containing 0.5 mg/ml collagenase (Type II; Worthington, Lakewood, NJ, USA) and 0.1 mg/ml protease (type XIV; Sigma, St. Louis, MO, USA) for 10 min. Ca^2+^-free Tyrode’s solution containing (in mM) 136 NaCl, 5.4 KCl, 0.33 NaH_2_PO_4_, 1 MgCl_2_, 10 glucose, and 10 HEPES (pH 7.4, adjusted with NaOH). The enzyme solution was then washed out, and the hearts were removed from the perfusion equipment. Left ventricles were placed in Petri dishes and were smoothly teased apart with forceps. Then, the cardiomyocytes were filtered through nylon mesh. The Ca^2+^ concentration was gradually increased to 1.0 mM, and the cells were stored at room temperature and used within 8 hours. Only cells from the left ventricular wall were used.

### Electrophysiology

Cardiomyocytes were patch-clamped in the whole-cell configuration of the patch-clamp technique in the current-clamp or the voltage-clamp mode. To record action potentials (APs), patch pipettes (2–5 MΩ) were loaded with an internal solution containing (in mM) 110 K^+^-aspartate, 30 KCl, 5 NaCl, 10 HEPES, 0.1 EGTA, 5 Mg-ATP, 5 Na_2_-creatine phosphates (pH 7.2, adjusted with KOH). The myocytes were superfused with normal Tyrode’s (NT) solution containing (in mM) 136 NaCl, 5.4 KCl, 0.33 NaH_2_PO_4_, 1 CaCl_2_, 1 MgCl_2_, 10 glucose, and 10 HEPES (pH 7.4, adjusted with NaOH). APs were obtained with 2-ms, 2- to 4-nA square pulses at multiple pacing cycle intervals (PCLs). We quantified triggered activity and changes in the resting membrane potential induced by Iso between 5 and 10 minutes after stimulation.

The Gap19 peptide (232ng/μL) and Cx43 antibody (2.5ng/μL) were added to the pipette solution to block Cx43 hemichannel activity. We used 2 or 3 cardiomyocytes per isolated heart per condition.

The two electrode-voltage clamp (TEVC) technique in *Xenopus* oocytes was used to test hemichannel currents from homomeric channels formed by Cx43 and Cx43S3A. Connexin clones were purchased from Origene (Rockville, MD, USA). Nhe1-linearized hCx43 and hCxCx43S3A were transcribed in vitro to cRNAs using the T7 Message Machine kit (Ambion, Austin, TX, USA). Electrophysiological data were collected using the Pclamp10 software. All recordings were made at room temperature (20-22°C). For Cx43 expressing oocytes, the recording solutions contained (in mM) 117 TEA and 5 HEPES and the extracellular Ca^2+^ concentration was 0.2 mM (pH 7.4, adjusted with N-Methyl-D-glucamine). Currents from oocytes were recorded 2 days after cRNA injection, using a Warner OC-725 amplifier (Warner Instruments, USA). Currents were sampled at 2 kHz and low pass filtered as 0.5 kHz. Microelectrode resistances were between 0.1 and 1.2 MΩ when filled with 3M KCl. Antisense oligonucleotides against Cx38 were injected to each oocyte to reduce the expression of endogenous Cx38 at 4 h after harvesting the oocytes (1mg/ml; using the sequence from Ebihara(Ebihara, 1996)). We assessed hemichannel currents and changes in the resting membrane potential evoked by NO at 10 minutes after stimulation. We used at least three oocytes from each independent frog.

### Telemetry Device Implantation

A Data Sciences International (DSI) telemetry transmitter, used to perform electrocardiography, was inserted as follows: After the animal was anesthetized by isoflurane, shaved, and aseptically prepared, a 1-2 cm transverse incision was made in the left inguinal area. A subcutaneous pocket was created on the left side of the abdomen up toward the caudal edge of the rib cage. The telemetric transmitter, which is a tubular disk 1 cm in diameter X 2 cm in length, was inserted into the subcutaneous pocket and secured to the abdominal wall with sutures in three places on the left flank. Two ECG lead wires were tunneled subcutaneously; the ends were stripped to provide transmission and then coiled and secured to the underlying muscle tissue. The positive ECG lead wire was tunneled to the lower left rib cage and fixed to the abdominal muscle; the negative ECG lead was tunneled to the caudal end of the right scapula and fixed to the pectoralis muscle on the right side. Finally, the transverse inguinal incision was closed in two layers using interrupted mattress sutures and 3-0 nylon. This procedure does not require externalization of the wires, thus minimizing the possibility of infection and allowing the animal to be monitored without interference.

### β-Adrenergic stress test Recording

After recovered from telemetry device implantation surgery (3-5 days), mice were subjected to a 24-hour Iso stress study. Mice were first weighed and separated into single cages that were placed on telemetry receivers. A 1-hour baseline reading was taken to monitor activity and obtain ECG readings. Next, mice were intraperitoneally injected with Iso (Isoproterenol, Sigma I6504, 5mg/kg). Mice were constantly recorded and observed for changes in activity and morbidity for 24 hours. Electrocardiographic data was analyzed by Lab Chart 8 software (Life Science Data Acquisition Software). Arrhythmias were scored based on a point system where: 0= no arrhythmias; 1= single premature ventricular contractions (PVCs); 2= double PVCs; 3= Triple PVCs or non-sustained ventricular tachycardia (VT); 4= sustained VT or atrioventricular (AV) block; and 5= death. We used between 6 and 7 independent mice per experiment.

### Dye perfusion and uptake in isolated hearts

Mice were heparinized (5000 Units/kg) and then anesthetized with isoflurane. Subsequently, mice were injected with either saline or Iso (5 mg/kg, IP). Twenty minutes following Iso or saline injection, mice were sacrificed, and hearts were extracted and cannulated in a Langendorff perfusion system. Hearts were perfused with Normal Tyrode’s buffer (NT) [in mM: 136 NaCl, 5.4 KCl, 0.33 NaH_2_PO_4_, 1 MgCl_2_, 1 CaCl_2_, 10 HEPES and 10 Glucose] at 37 °C degrees for 10 minutes, followed by NT containing ethidium bromide (5 μM) or propidium iodide (50 μM) for 20 minutes and then NT buffer for 5 minutes to wash out the dye. Hearts were fixed overnight in 4% paraformaldehyde (Sigma, St. Louis, MO, USA), placed into 30% sucrose solution in PBS (Sigma, St. Louis, MO, USA) for 12 hours, then embedded in O.C.T (Tissue-Tek, USA). Afterwards, 10 μm cryosections were made, slides were thawed to room temperature, washed in PBS, and Alexa Fluor Wheat Germ Agglutinin 555 (Invitrogen, NY, USA) was applied for 20 minutes. Slides were then washed in PBS and mounted with mounting reagent containing DAPI (Invitrogen, NY, USA). Slides were imaged using a 200 Axiovert fluorescence microscope (Zeiss, Oberkochen, Germany). To calculate ethidium fluorescence in ImageJ, DAPI stained nuclei were identified, created as ROI and individual nuclei (100-150 per image) mean fluorescent intensities were measured. Then, the ROI outlines were projected onto the corresponding ethidium image, where individual fluorescent intensities were measured, capturing ethidium signal within all nuclei. Ethidium intensity was then divided by DAPI nuclei intensity for each respective ROI signal, then the mean ratio was calculated for all nuclei in the image. Three images per heart were evaluated in a blinded manner.

### Biotin Perfusion of Isolated Hearts

Mice were heparinized and then anesthetized with isoflurane. Once unconscious, mice were sacrificed, and hearts were extracted and cannulated in a Langedorff perfusion system. Hearts were perfused with NT for 5 minutes, changed to NT buffer plus Biotin (EZ-Link NHS Biotin, 0.5 mg/mL, Thermo Scientific, Waltham, MA, USA) for 60 minutes (0.25 ml/min flow rate), and washed out for 10 minutes with NT buffer plus 15 mM Glycine. Left ventricular tissue was later homogenized in HEN buffer (in mM: 250 HEPES, 1 EDTA, 0.1 Neocuproine, pH 7.7) with 2x HALT protease inhibitors (Thermo Scientific, NY, USA) and then centrifuged at 16,000 g for 10 minutes. Following protein concentration determination, 50 μl of streptavidin beads (Thermo Scientific, NY, USA) were added to 200 μg protein and nutated for 90 minutes at 4^0^C with vortexing every 10 minutes. Samples were then centrifuged at 16,000 g for 2 minutes and the supernatant was dropped.

Next, the streptavidin pellet was resuspended in fresh lysis buffer containing 0.1% Triton X-100 and centrifuged for 1 minute at 16,000 g. The pellet was then washed with PBS (pH 7.4) and centrifuged. 25μl of 2x Laemmli sample buffer was added and heated at 100 ^0^C for 5 minutes to disrupt the biotin-streptavidin interaction. The heated samples were then centrifuged for 1 minute at 16,000 g and the supernatant was run along with total protein extracts without streptavidin pulldown on SDS-PAGE.

### Tissue Immunofluorescence and Cx43 Quantification

Immunofluorescent staining was performed on cryo-embedded cardiac sections with Cx43 (Sigma C6219, 1:2000, rabbit) and N-Cadherin (Invitrogen 33-3900, 1:400, mouse) antibodies as described in^19^. Confocal Z-stacks at a step size of 0.5μm thickness (approximately 12 slices per image) were acquired at 60x magnification on an Olympus Fluoview 1000 Confocal Laser Scanning Microscope using the Fluoview software. The Cx43/N-Cadherin images were then separated into their separate channels and processed as maximum intensity z-stack projections in Fiji prior to analysis. Maximum intensity z-stacks were processed and Cx43 intensities at N-Cadherin positive intercalated discs were calculated as described in^19^.

### Western Blotting

Protein samples from the left ventricular heart wall or from injected *Xenopus* oocytes were separated by 10% SDS-PAGE and transferred onto a PVDF membrane (BioRad, Hercules, CA, USA). The primary, Cx43 (Sigma St. Louis, MO, USA, #C8093, 1:2000, mouse), and secondary (Pierce, Rockford, IL, USA; 1/5000). Protein bands were detected with the SuperSignal® West Femto (Pierce, Rockford, IL, USA). Molecular mass was estimated with pre-stained markers (BioRad, Hercules, CA, USA). Protein bands were analyzed using the ImageJ software (NIH, USA).

### Detection of S-nitrosylated proteins

S-nitrosylated proteins were isolated from either mouse heart ventricular samples or *Xenopus* oocytes expressing Cx43 WT or Cx43SA. Heart tissue or *Xenopus* oocytes were homogenized in HEN buffer (in mM: 250 HEPES, 1 EDTA, 0.1 Neucoproine, pH 7.7) containing protease inhibitors. Samples containing 200 μg protein were treated by the biotin-switch method to pull down all S-nitrosylated proteins as previously described(Lillo et al., 2019). S-nitrosylated proteins were separated using 10% SDS-PAGE and transferred onto a PVDF membrane (BioRad, Hercules, CA, USA). A monoclonal anti-Cx43 antibody (Sigma, St. Louis, MO, USA, #C8093, 1:2000, mouse) was used to detect Cx43 protein. For all Western blot analyses, the intensity of the signal was evaluated using ImageJ (NIH, USA). We used 6 independent hearts per treatment.

### Analysis of protein-to-protein association

The subcellular distribution and possible spatial association between S-NO and Cx43 were evaluated by Proximity Ligation Assay(Soderberg et al., 2006) (Sigma, St. Louis, MO, USA). Tissue sections (6 μm) were blocked and incubated with two primary antibodies from different species, which were then detected using oligonucleotide-conjugated secondary antibodies as described in the manufacturer’s protocols. Monoclonal anti-Cx43 (Sigma, St. Louis, MO, USA, #C8093, 1:200) and anti-S-nitrosocysteine (Sigma, St. Louis, MO, USA, #N5411, 1:100) antibodies were used. Images were visualized with a 200 Axiovert fluorescence microscope (Zeiss, Oberkochen, Germany). We used 5 independent hearts per treatment.

### Chemicals

HEPES, cAMP, Na2-creatine phosphate, K^+^-aspartate and N-Methyl-D-glucamine were purchased from Sigma-Aldrich (St. Louis, MO, USA). Isoproterenol was obtained from Sigma Aldrich (420355, USA) and collagenase type II from Worthington (Lakewood, NJ, USA). Gap19 was purchased from Tocris (Minneapolis, MN, USA).

### Statistical analysis

Values are displayed as mean ± standard error. Comparisons between groups were made using paired Student’s t-test, one-way ANOVA or two-way ANOVA plus Tukey post-hoc test, as appropriate. P < 0.05 was considered significant. Each legend figure indicates the respective *n* value.

## Acknowledgements

We thank Dr. Glenn I. Fishman (New York University) for providing the original S3A founder mice for our colonies.

---

**Figure S1. S3A mice displayed arrhythmogenic behaviors during the dark cycle with and without isoproterenol (Iso). A)** Quantification of arrhythmogenic events (including PVC, double PVC, VT, or AV block) from 4-6-month-old mice during the dark and light cycle without Iso in WT, S3A and S3A mice treated with Gap19 via retroorbital injection (10 μg/kg). The number in parentheses indicates the *n* value. Comparisons between groups were made using two-way ANOVA plus Tukey post-hoc test. *<0.05 vs wild-type, †P<0.05 vs S3A. **B**) Quantification of arrhythmogenic events (including PVC, double PVC, VT, or AV block) from 4-6-month-old mice during the dark and light cycle upon Iso stimulation (5 mg/kg, IP) in WT, S3A and S3A mice treated with Gap19 via retroorbital injection (10 μg/kg). The number in parentheses indicates the *n* value. Comparisons between groups were made using two-way ANOVA plus Tukey post-hoc test. *<0.05 vs WT, †P<0.05 vs S3A, **<0.05 vs S3A Dark Cycle. **C**) Movements detected using an *in vivo* telemetry system in WT, S3A and S3A mice treated with Gap19 via retroorbital injection (10 μg/kg). Arrow indicates Iso stimulation. Comparisons between groups during the total behavior recording were made using two-way ANOVA plus Tukey post-hoc test. *<0.05 vs WT.

## REFERENCES

Abudara, V., J. Bechberger, M. Freitas-Andrade, M. De Bock, N. Wang, G. Bultynck, C.C. Naus, L. Leybaert, and C. Giaume. 2014. The connexin43 mimetic peptide Gap19 inhibits hemichannels without altering gap junctional communication in astrocytes. Front Cell Neurosci. 8:306.

Bruce, A.F., S. Rothery, E. Dupont, and N.J. Severs. 2008. Gap junction remodelling in human heart failure is associated with increased interaction of connexin43 with ZO-1. Cardiovasc Res. 77:757–765.

Contreras, J.E., J.C. Saez, F.F. Bukauskas, and M.V. Bennett. 2003. Gating and regulation of connexin 43 (Cx43) hemichannels. Proc Natl Acad Sci U S A. 100:11388–11393.

De Smet, M.A., A. Lissoni, T. Nezlobinsky, N. Wang, E. Dries, M. Perez-Hernandez, X. Lin, M. Amoni, T. Vervliet, K. Witschas, E. Rothenberg, G. Bultynck, R. Schulz, A.V. Panfilov, M. Delmar, K.R. Sipido, and L. Leybaert. 2021. Cx43 hemichannel microdomain signaling at the intercalated disc enhances cardiac excitability. J Clin Invest. 131.

Ebihara, L. 1996. Xenopus connexin38 forms hemi-gap-junctional channels in the nonjunctional plasma membrane of Xenopus oocytes. Biophys J. 71:742–748.

Fauconnier, J., J. Thireau, S. Reiken, C. Cassan, S. Richard, S. Matecki, A.R. Marks, and A. Lacampagne. 2010. Leaky RyR2 trigger ventricular arrhythmias in Duchenne muscular dystrophy. Proc Natl Acad Sci U S A. 107:1559–1564.

Figueroa, X.F., M.A. Lillo, P.S. Gaete, M.A. Riquelme, and J.C. Saez. 2013. Diffusion of nitric oxide across cell membranes of the vascular wall requires specific connexin-based channels. Neuropharmacology. 75:471–478.

Forrester, M.T., M.W. Foster, M. Benhar, and J.S. Stamler. 2009. Detection of protein S-nitrosylation with the biotin-switch technique. Free Radic Biol Med. 46:119–126.

Ghazizadeh, Z., T. Kiviniemi, S. Olafsson, D. Plotnick, M.E. Beerens, K. Zhang, L. Gillon, M.J. Steinbaugh, V. Barrera, and S.H.J.C. Sui. 2020. Metastable atrial state underlies the primary genetic substrate for MYL4 mutation-associated atrial fibrillation. 141:301–312.

Gollob, M.H. 2006. Cardiac connexins as candidate genes for idiopathic atrial fibrillation. Curr Opin Cardiol. 21:155–158.

Gonzalez, D.R., F. Beigi, A.V. Treuer, and J.M. Hare. 2007. Deficient ryanodine receptor S-nitrosylation increases sarcoplasmic reticulum calcium leak and arrhythmogenesis in cardiomyocytes. Proc Natl Acad Sci U S A. 104:20612–20617.

Gonzalez, J.P., J. Ramachandran, E. Himelman, M.A. Badr, C. Kang, J. Nouet, N. Fefelova, L.H. Xie, N. Shirokova, J.E. Contreras, and D. Fraidenraich. 2018. Normalization of connexin 43 protein levels prevents cellular and functional signs of dystrophic cardiomyopathy in mice. Neuromuscul Disord. 28:361–372.

Gonzalez, J.P., J. Ramachandran, L.H. Xie, J.E. Contreras, and D. Fraidenraich. 2015. Selective Connexin43 Inhibition Prevents Isoproterenol-Induced Arrhythmias and Lethality in Muscular Dystrophy Mice. Sci Rep. 5:13490.

Grimm, M., and J.H. Brown. 2010. Beta-adrenergic receptor signaling in the heart: role of CaMKII. J Mol Cell Cardiol. 48:322–330.

Gutstein, D.E., G.E. Morley, H. Tamaddon, D. Vaidya, M.D. Schneider, J. Chen, K.R. Chien, H. Stuhlmann, and G.I. Fishman. 2001. Conduction slowing and sudden arrhythmic death in mice with cardiac-restricted inactivation of connexin43. Circ Res. 88:333–339.

Hegyi, B., S. Morotti, C. Liu, K.S. Ginsburg, J. Bossuyt, L. Belardinelli, L.T. Izu, Y. Chen-Izu, T. Banyasz, E. Grandi, and D.M. Bers. 2019. Enhanced Depolarization Drive in Failing Rabbit Ventricular Myocytes: Calcium-Dependent and beta-Adrenergic Effects on Late Sodium, L-Type Calcium, and Sodium-Calcium Exchange Currents. Circ Arrhythm Electrophysiol. 12:e007061.

Himelman, E., M.A. Lillo, J. Nouet, J.P. Gonzalez, Q. Zhao, L.H. Xie, H. Li, T. Liu, X.H. Wehrens, P.D. Lampe, G.I. Fishman, N. Shirokova, J.E. Contreras, and D. Fraidenraich. 2020. Prevention of connexin-43 remodeling protects against Duchenne muscular dystrophy cardiomyopathy. J Clin Invest. 130:1713–1727.

Huang, R.Y., J.G. Laing, E.M. Kanter, V.M. Berthoud, M. Bao, H.W. Rohrs, R.R. Townsend, and K.A. Yamada. 2011. Identification of CaMKII phosphorylation sites in Connexin43 by high-resolution mass spectrometry. J Proteome Res. 10:1098–1109.

Johnson, R.G., H.C. Le, K. Evenson, S.W. Loberg, T.M. Myslajek, A. Prabhu, A.M. Manley, C. O’Shea, H. Grunenwald, M. Haddican, P.M. Fitzgerald, T. Robinson, B.A. Cisterna, J.C. Saez, T.F. Liu, D.W. Laird, and J.D. Sheridan. 2016. Connexin Hemichannels: Methods for Dye Uptake and Leakage. The Journal of membrane biology. 249:713–741.

Kim, J.C., M. Perez-Hernandez, F.J. Alvarado, S.R. Maurya, J. Montnach, Y. Yin, M. Zhang, X. Lin, C. Vasquez, Heguy, F.X. Liang, S.H. Woo, G.E. Morley, E. Rothenberg, A. Lundby, H.H. Valdivia, M. Cerrone, and M. Delmar. 2019b. Disruption of Ca(2+)i Homeostasis and Connexin 43 Hemichannel Function in the Right Ventricle Precedes Overt Arrhythmogenic Cardiomyopathy in Plakophilin-2-Deficient Mice. Circulation. 140:1015–1030.

Kleber, A.G., and J.E. Saffitz. 2014. Role of the intercalated disc in cardiac propagation and arrhythmogenesis. Frontiers in physiology. 5:404.

Kyrychenko, S., E. Polakova, C. Kang, K. Pocsai, N.D. Ullrich, E. Niggli, and N. Shirokova. 2013. Hierarchical accumulation of RyR post-translational modifications drives disease progression in dystrophic cardiomyopathy. Cardiovasc Res. 97:666–675.

Lillo, M.A., and J.E. Contreras. 2021. Opening the floodgates: An emerging role for Connexin-43 hemichannels in the heart. Cell Calcium. 97:102410.

Lillo, M.A., E. Himelman, N. Shirokova, L.H. Xie, D. Fraidenraich, and J.E. Contreras. 2019. S-nitrosylation of connexin43 hemichannels elicits cardiac stress-induced arrhythmias in Duchenne muscular dystrophy mice. JCI Insight. 4.

Lima, B., M.T. Forrester, D.T. Hess, and J.S. Stamler. 2010. S-nitrosylation in cardiovascular signaling. Circ Res. 106:633–646.

Lissoni, A., P. Hulpiau, T. Martins-Marques, N. Wang, G. Bultynck, R. Schulz, K. Witschas, H. Girao, M. De Smet, and L. Leybaert. 2019. RyR2 regulates Cx43 hemichannel intracellular Ca2+-dependent activation in cardiomyocytes. Cardiovasc Res.

Lissoni, A., P. Hulpiau, T. Martins-Marques, N. Wang, G. Bultynck, R. Schulz, K. Witschas, H. Girao, M. De Smet, and L. Leybaert. 2021. RyR2 regulates Cx43 hemichannel intracellular Ca2+-dependent activation in cardiomyocytes. Cardiovasc Res. 117:123–136.

Mani, S.K., E.A. Egan, B.K. Addy, M. Grimm, H. Kasiganesan, T. Thiyagarajan, L. Renaud, J.H. Brown, C.B. Kern, and D.R. Menick. 2010. beta-Adrenergic receptor stimulated Ncx1 upregulation is mediated via a CaMKII/AP-1 signaling pathway in adult cardiomyocytes. J Mol Cell Cardiol. 48:342–351.

Marx, S.O., S. Reiken, Y. Hisamatsu, T. Jayaraman, D. Burkhoff, N. Rosemblit, and A.R. Marks. 2000. PKA phosphorylation dissociates FKBP12.6 from the calcium release channel (ryanodine receptor): defective regulation in failing hearts. Cell. 101:365–376.

Oxford, E.M., H. Musa, K. Maass, W. Coombs, S.M. Taffet, and M. Delmar. 2007. Connexin43 remodeling caused by inhibition of plakophilin-2 expression in cardiac cells. Circ Res. 101:703–711.

Pfeiffer, S., E. Leopold, K. Schmidt, F. Brunner, and B. Mayer. 1996. Inhibition of nitric oxide synthesis by NG-nitro-L-arginine methyl ester (L-NAME): requirement for bioactivation to the free acid, NG-nitro-L-arginine. British journal of pharmacology. 118:1433–1440.

Qu, J., F.M. Volpicelli, L.I. Garcia, N. Sandeep, J. Zhang, L. Marquez-Rosado, P.D. Lampe, and G.I. Fishman. 2009. Gap junction remodeling and spironolactone-dependent reverse remodeling in the hypertrophied heart. Circ Res. 104:365–371.

Remo, B.F., S. Giovannone, and G.I. Fishman. 2012. Connexin43 cardiac gap junction remodeling: lessons from genetically engineered murine models. J Membr Biol. 245:275–281.

Remo, B.F., J. Qu, F.M. Volpicelli, S. Giovannone, D. Shin, J. Lader, F.Y. Liu, J. Zhang, D.S. Lent, G.E. Morley, and G.I. Fishman. 2011. Phosphatase-resistant gap junctions inhibit pathological remodeling and prevent arrhythmias. Circ Res. 108:1459–1466.

Saffitz, J.E. 2009. Desmosome mutations in arrhythmogenic right ventricular cardiomyopathy: important insight but only part of the picture. Circ Cardiovasc Genet. 2:415–417.

Saffitz, J.E., A. Asimaki, and H. Huang. 2009. Arrhythmogenic right ventricular cardiomyopathy: new insights into disease mechanisms and diagnosis. J Investig Med. 57:861–864.

Severs, N.J., S.R. Coppen, E. Dupont, H.I. Yeh, Y.S. Ko, and T. Matsushita. 2004a. Gap junction alterations in human cardiac disease. Cardiovasc Res. 62:368–377.

Severs, N.J., E. Dupont, S.R. Coppen, D. Halliday, E. Inett, D. Baylis, and S. Rothery. 2004b. Remodelling of gap junctions and connexin expression in heart disease. Biochimica et biophysica acta. 1662:138–148.

Severs, N.J., E. Dupont, N. Thomas, R. Kaba, S. Rothery, R. Jain, K. Sharpey, and C.H. Fry. 2006. Alterations in cardiac connexin expression in cardiomyopathies. Adv Cardiol. 42:228–242.

Soderberg, O., M. Gullberg, M. Jarvius, K. Ridderstrale, K.J. Leuchowius, J. Jarvius, K. Wester, P. Hydbring, F. Bahram, L.G. Larsson, and U. Landegren. 2006. Direct observation of individual endogenous protein complexes in situ by proximity ligation. Nat Methods. 3:995–1000.

Thibodeau, I.L., J. Xu, Q. Li, G. Liu, K. Lam, J.P. Veinot, D.H. Birnie, D.L. Jones, A.D. Krahn, R. Lemery, B.J. Nicholson, and M.H. Gollob. 2010. Paradigm of genetic mosaicism and lone atrial fibrillation: physiological characterization of a connexin 43-deletion mutant identified from atrial tissue. Circulation. 122:236–244.

Tse, G.J.J.o.a. 2016. Mechanisms of cardiac arrhythmias. 32:75–81.

Vielma, A.Z., M.P. Boric, and D.R. Gonzalez. 2020. Apocynin Treatment Prevents Cardiac Connexin 43 Hemichannels Hyperactivity by Reducing Nitroso-Redox Stress in Mdx Mice. Int J Mol Sci. 21.

Vielma, A.Z., L. Leon, I.C. Fernandez, D.R. Gonzalez, and M.P. Boric. 2016. Nitric Oxide Synthase 1 Modulates Basal and beta-Adrenergic-Stimulated Contractility by Rapid and Reversible Redox-Dependent S-Nitrosylation of the Heart. PLoS One. 11:e0160813.

Wang, N., E. De Vuyst, R. Ponsaerts, K. Boengler, N. Palacios-Prado, J. Wauman, C.P. Lai, M. De Bock, E. Decrock, M. Bol, M. Vinken, V. Rogiers, J. Tavernier, W.H. Evans, C.C. Naus, F.F. Bukauskas, K.R. Sipido, G. Heusch, R. Schulz, G. Bultynck, and L. Leybaert. 2013. Selective inhibition of Cx43 hemichannels by Gap19 and its impact on myocardial ischemia/reperfusion injury. Basic Res Cardiol. 108:309.

Zhu, W.-Z., S.-Q. Wang, K. Chakir, D. Yang, T. Zhang, J.H. Brown, E. Devic, B.K. Kobilka, H. Cheng, and R.-P.J.T.J.o.c.i. Xiao. 2003. Linkage of β 1-adrenergic stimulation to apoptotic heart cell death through protein kinase A–independent activation of Ca 2+/calmodulin kinase II. 111:617–625.

