## Supplementary figures and images for "Remodeled Connexin 43 hemichannels alter cardiac excitability and promote arrhythmias"

### Figure S1

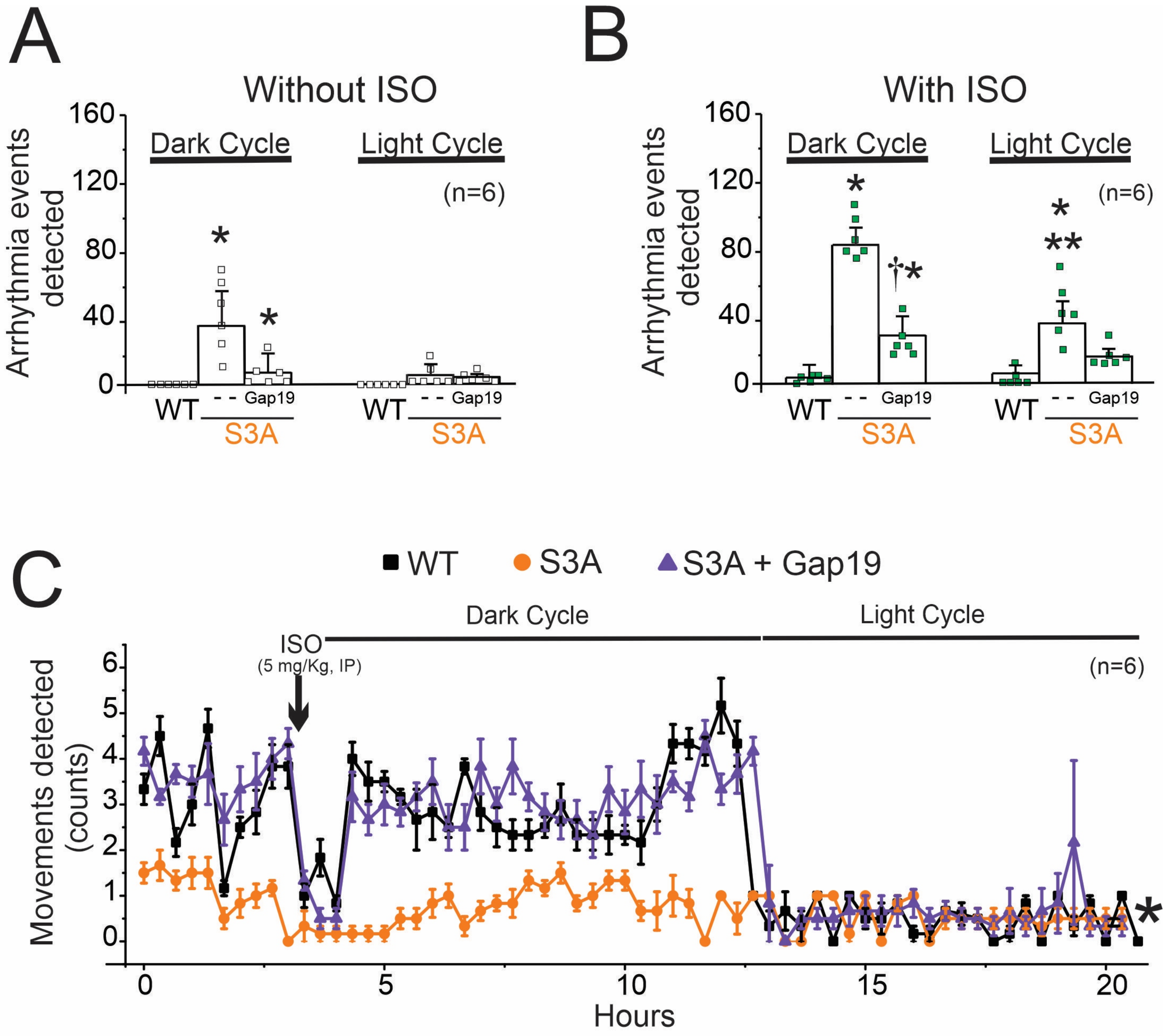
